# Unveiling the neural dynamics of conscious perception in rapid object recognition

**DOI:** 10.1101/2023.11.08.566069

**Authors:** Saba Charmi Motlagh, Marc Joanisse, Boyu Wang, Yalda Mohsenzadeh

## Abstract

Our brain excels at recognizing objects, even when they flash by in a rapid sequence. However, the neural processes determining whether a target image in a rapid sequence can be recognized or not remained elusive. We used electroencephalography (EEG) to investigate the temporal dynamics of brain processes that shape perceptual outcomes in these challenging viewing conditions. Using naturalistic images and advanced multivariate pattern analysis (MVPA) techniques, we probed the brain dynamics governing conscious object recognition. Our results show that although initially similar, the processes for when an object can or cannot be recognized diverge around 180ms post-appearance, coinciding with feedback neural processes. Decoding analyses indicate that object categorization can occur at ∼120ms through feedforward mechanisms. In contrast, object identification is resolved at ∼190ms after target onset, suggesting involvement of recurrent processing. These findings underscore the importance of recurrent neural connections in object recognition and awareness in rapid visual presentations.

## Introduction

Every day, from the first glimpse of morning light to the hazy twilight hours, our eyes witness countless objects. Scattered items on the tables, dogs wagging their tails as they walk alongside their owners, and people traversing from one room to another— these are a few snapshots from our daily visual experience. The human brain effortlessly sifts through countless possibilities to rapidly recognize these objects^1,2^. It is crucial for our survival to perceive the external world around us quickly and accurately; however, unlike the controlled research settings where objects are presented in isolation, our daily lives are filled with a multitude of objects existing in cluttered spaces^3,4^. As a result, the process of object recognition faces additional challenges in various conditions, including object deletion, occlusion, and crowding^5–7^.

One of these challenges is limited presentation time. Often, in daily life, we only see objects for a short period of time. For example, while we are driving or running, objects often move quickly and are in our visual field for a limited time. In laboratory settings, rapid serial visual presentation (RSVP) is one of the most effective experimental designs used to study this challenging condition^8–11^. RSVP involves rapidly presented sequence of images embedded with one or several targets. The temporal constraints of recognizing a target within an RSVP stream may be attributed to the interference caused by processing multiple images in rapid succession. If a new image arrives at a rate that exceeds the visual system’s temporal capabilities, target images may fail to be perceived^10,12^. Numerous studies have investigated the intricacies of object recognition within the RSVP paradigm^13–17^; however, the precise neural processes that determine whether a target image in RSVP is recognized or not are still unclear.

Hierarchical processing serves as a fundamental framework for understanding visual perception. After an image is presented, a cascade of feedforward connections activates the successive hierarchical levels of the visual cortex. The activation spreads from lower-level to higher-level areas within the visual hierarchy. A growing body of literature suggests that this bottom-up activation of visual areas enables the visual system to build an initial coarse visual representation, and is sufficient for detection and rapid categorization tasks^18–24^. While feedforward connections transmit information from lower to higher visual areas, horizontal and feedback connections provide input from cells at the same level and from higher levels in the visual hierarchy, respectively. This recurrent processing might be critical for solving certain visual tasks such as visual occlusion or figure-ground segregation, which require integration of information from outside the receptive field^24–27^ and it can be distinguished from the initial feedforward sweep based on its processing time in visual tasks. According to primate electrophysiological studies^28–31^, the fast feedforward sweep is completed within approximately 100-150 ms after the visual onset. As a result, visually-evoked neural responses beyond these latencies can thus be attributed to recurrent processing^5,14,18,24,32^.

Neuroimaging and computational findings have highlighted the presence and contribution of recurrent processes to object recognition, specifically during challenging viewing conditions^5,14,33^. Additionally, studies that have contrasted correctly and incorrectly recognized stimuli suggested that they have similar temporal dynamics during the early stages of processing and diverge at later stages^34–36^. Nonetheless, the underlying cause of this divergence remains a topic of ongoing debate.

Beyond categorization, our brain can uniquely identify individual objects in a class. To correctly identify an object, we need to ignore changes in size, position, lighting, and viewpoint, but tolerate some changes in shape, although other differences in shape can imply different identities. On the other hand, our brain can tell what class of object something belongs to. In the context of identification, it is necessary to differentiate between objects that may share physical similarities. However, when it comes to categorization, we need to draw broader conclusions that encompass objects with varying physical attributes. The degree of this broader categorization depends on the level of abstraction within the categories (objects can be categorized at several levels of abstraction, for example, animal, mammal, cat, Abyssinian, Max)^37^. Several studies have suggested that the human brain can get the “gist” of an object in a shorter time in comparison to identifying it at a finer grain^38–40^. But what accounts for this disparity in the timing of categorization and identification?

In the present study, we aim to investigate the temporal dynamics of neural processes that determine whether a target image in RSVP is correctly (visible) or incorrectly (invisible) recognized and to elucidate the contributions of feedforward and recurrent processing in shaping these perceptual outcomes. Moreover, we wanted to illuminate the neural processes contributing to target identification and perceptual categorization. For this, we use naturalistic target images and multivariate pattern analysis (MVPA) to directly relate behavior and electrophysiological measurements. The advantage of using naturalistic images is to replicate the complexity of real-world scenarios^41^, which might improve the generality of our findings. Recent research efforts have applied MPVA approaches such as pattern classification and representational similarity analysis (RSA)^42^ to demonstrate that magnetoencephalography (MEG) and EEG signals contain information about a wide range of sensory and cognitive processes^5,13,14,43–47^. Here, we use this approach to examine whether visible and invisible conditions exhibit comparable temporal dynamics during the early stages of visual processing in line with previous ERP studies^34,48^, but different temporal dynamics during the later stages of visual processing. In addition, MVPA is employed to investigate the timing of perceptual categorization and object identification in the human brain and whether categorization precedes identification. Such results would suggest that recurrent processes play an important role in bringing object details into awareness, especially in challenging conditions.

## Results

We collected EEG data from 30 participants while they watched a rapid series of images each presented for 17 ms. Each trial was composed of 11 images with the middle image as the target and the other 10 images as masks. Targets were chosen to depict people (including images of faces and body parts) and objects (including images of animals and non-animal objects) and masks were scrambled images generated using our target images (see methods section; Figure 1). Participants performed a forced choice task in which they had to choose the target image among four options on the screen. Participants’ performance was significantly above chance (*p* < 0.0001 for all subcategories of images; Figure 2) in spite of the difficulty of the task.

**Figure 1.**
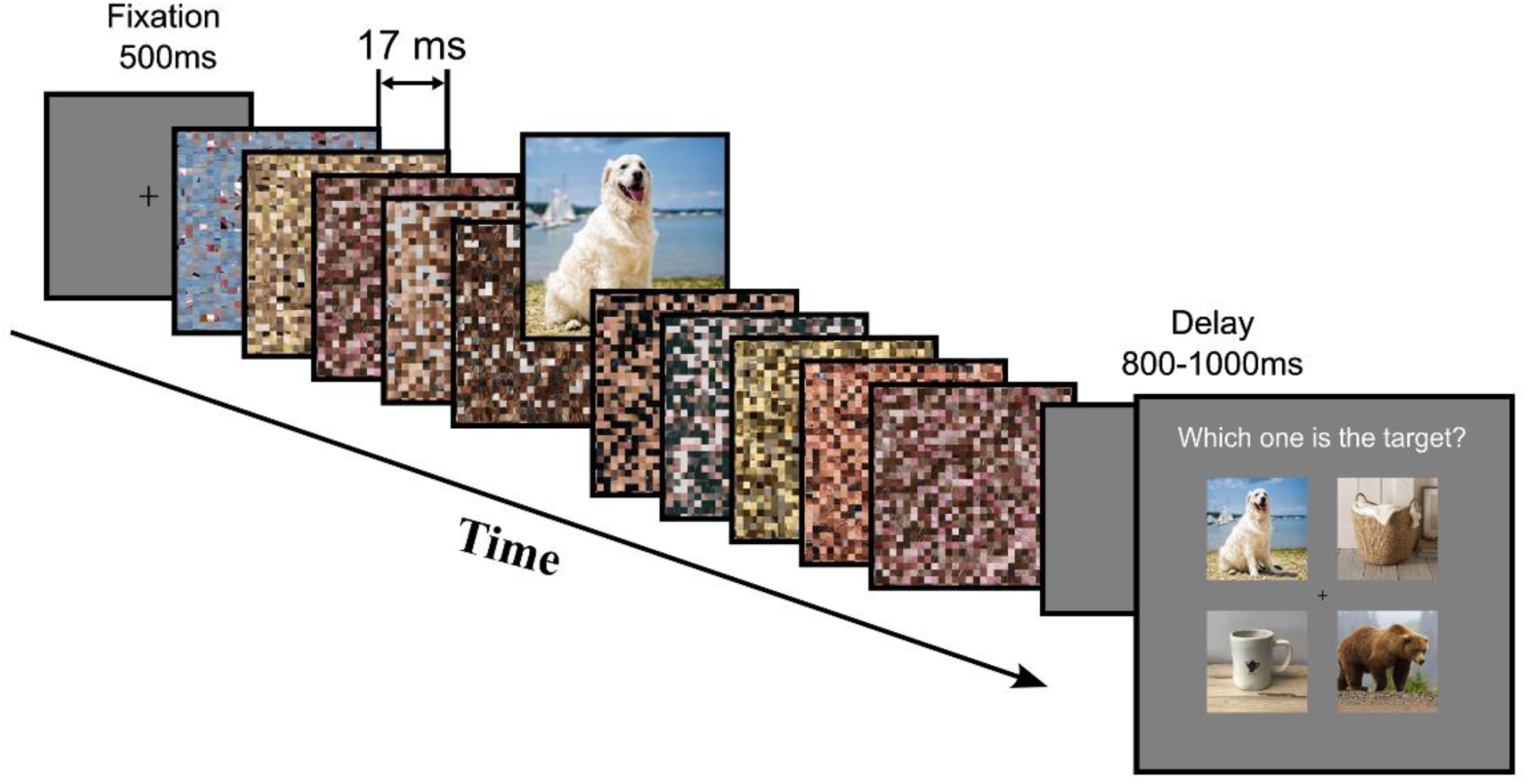
| Experimental procedure. Experimental procedure. Each trial starts with 500 ms of fixation followed by 11 images with the middle (sixth) image as the target. Other images in the stream are scrambled images and the target is chosen from either the people category or objects category. The time interval between two consecutive images is 17 ms. At the end of each trial, after a short delay of 800-1000 ms, participants were asked to report the target by choosing it among four options.

**Figure 2.**
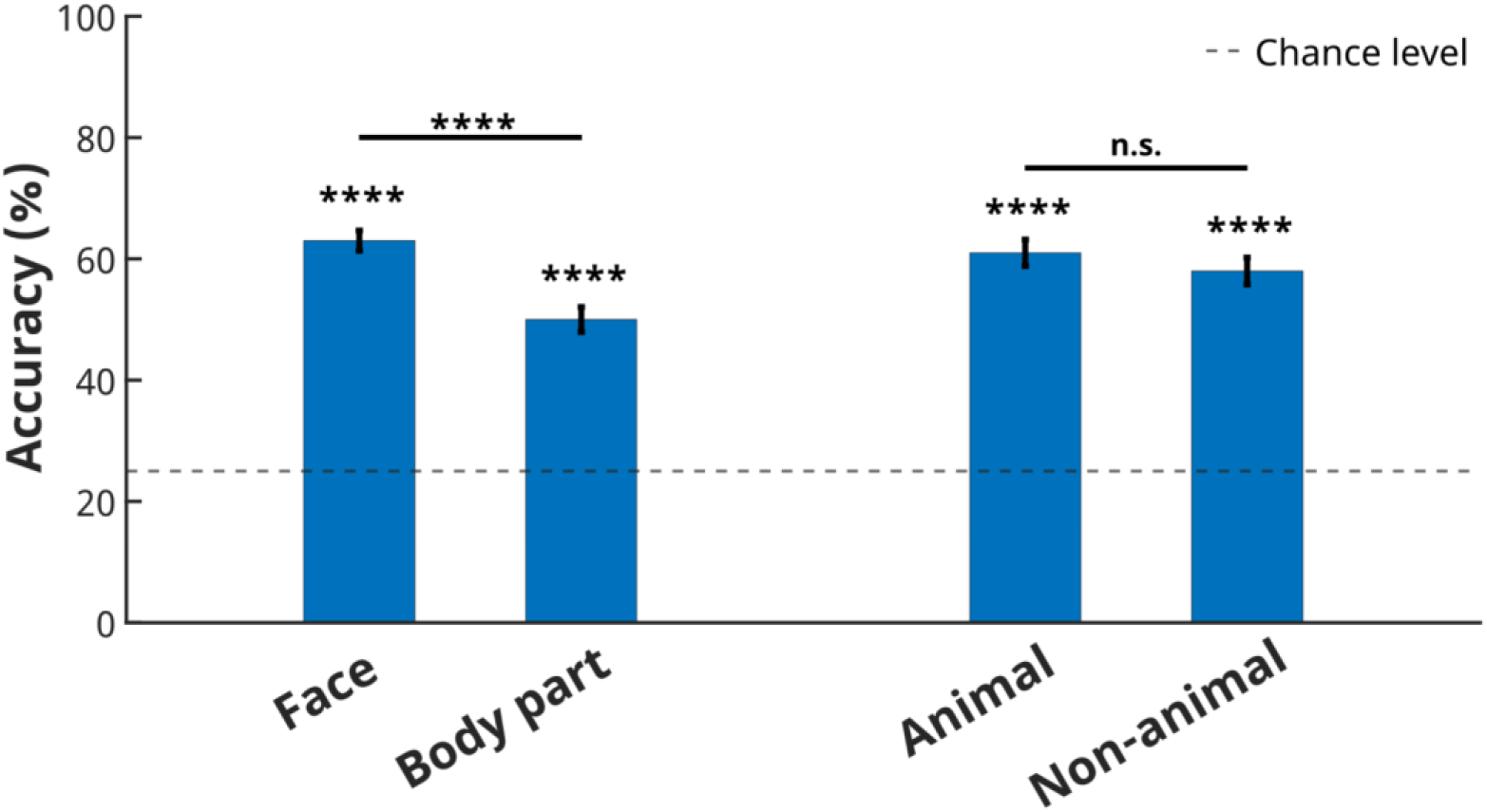
| Behavioral results. Average behavioral performance (N=30) in target image identification for people (divided into face and body part subgroups) and objects (divided into animal and non-animal subgroups). The dashed line indicates the chance level performance (25%). Participants performed well above chance for all four subgroups of images. Error bars represent SEM.

### 1. Visual recognition in challenging visibility conditions demands more than feedforward processing

To test how and when target images are discriminated in the human brain, we performed multivariate pattern analysis of the EEG data. In this analysis, we used a linear Support Vector Machine (SVM) and leave-one-out cross-validation to classify the trials for each pair of target images, yielding decoding accuracy of the classifier in separating images in each pair. Using the decoding accuracies for all possible pairs of images, we formed a 40×40 Representational Dissimilarity Matrix (RDM), at each time point. We averaged over all elements in this matrix (excluding diagonal elements) to find the grand total decoding time series for each participant. The average decoding accuracy (grand total) time series across all participants is shown in Figure 3A. By tracking the changes in decoding accuracy over time, we can see that decoding accuracy rises significantly above chance 25 ms after target onset. There are two early salient peaks in this grand total curve: the first peak happens ∼100 ms after target onset and the second one ∼200 ms after stimulus onset. Previous studies have shown that the rapid feedforward propagation of information will be completed around 100-150 ms following the onset of visual stimulation^14,24,49^. The first peak in neural decoding aligns with the timeframe typically associated with feedforward processing. The appearance of the second peak suggests that there is more complex neural processing occurring after this initial feedforward phase.

**Figure 3.**
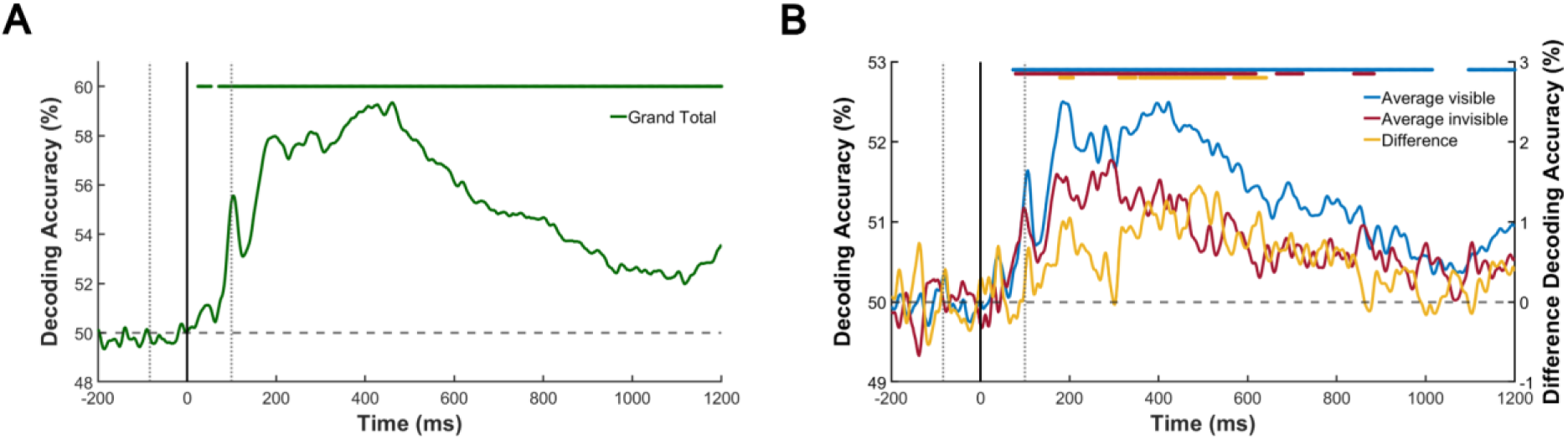
| Grand total, visible, and invisible curves. (A) Average grand total decoding accuracy for all participants. The color-coded line above the plot shows significant time points (N=30; significant time points were evaluated with one-sided sign permutation tests, cluster defining threshold *p* < 0.01, and corrected significance level *p* < 0.05). The dashed horizontal line indicates chance level performance. The dotted vertical lines show the RSVP onset/offset and the solid vertical line shows the target onset. (B) Average decoding accuracies for the visible and invisible conditions and the difference between the two. The left y-axis indicates decoding accuracy for visible and invisible conditions, and the right y-axis indicates the difference in decoding accuracy for visible and invisible conditions. The color-coded lines above the plots show significant time points for each condition (N=30; significant time points were evaluated with one-sided sign permutation tests, cluster defining threshold *p* < 0.01, and corrected significance level *p* < 0.05). The dashed horizontal line indicates chance level performance. The dotted vertical lines show the RSVP onset/offset and the solid vertical line shows the target onset.

### 2. Recurrent processing plays a vital role in the visibility of objects

Although target images can be resolved at the level of individual images using EEG data and MVPA, our behavioral data show that target images are invisible to our participants in a relatively large number of trials. What is the difference in temporal neural dynamics that leads to a target being visible or invisible? To address this question, we separated visible and invisible trials for each target image in each participant’s EEG data. Using MVPA, we formed two 40×40 RDMs for the visible and invisible trials separately, rows and columns in each indexed by target images. We constructed these matrices for each participant and each time point. The average decoding time series and the difference in decoding accuracy between visible and invisible trials are shown in Figure 3B. These time series show that the visible and invisible trials become significantly different from each other ∼180 ms after target onset and this difference peaks at ∼200 ms after target onset. The peak of the difference time series coincides with the second peak of target image decoding. This cannot be explained only by feedforward processing, as it occurs later than the first stage of processing. This late difference between visible and invisible cases is consistent with the timing of recurrent processing as shown in previous studies^18,49^.

### 3. The recognized targets benefit from more sustained neural processing

Various processing stages can occur at different time points. The neuronal activity associated with these processing stages can exhibit either persistence (a sustained pattern of activity over time) or transience (rapid changes over time). When neuronal activity persists over time, it is expected that EEG signals will remain similar during those intervals^45^. To investigate the persistence of neural responses to the visible and invisible trials, we used the temporal generalization approach^46^. For each participant, a linear SVM classifier was trained to pairwise decode EEG data for different target images on time point (t) and then tested using EEG data at another time point (t’) to investigate shared representations across time. The temporal generalization maps were then extracted for all pairs of time points (t,t’) and were averaged across all participants. The temporal generalization maps for visible and invisible cases and the difference between these two cases are shown in Figures 4A, 4B, and 4C. The white contours show the time points when the accuracies are significantly above chance (N=30, one-sided sign permutation tests, cluster-defining threshold *p* < 0.01, and corrected significance level *p* < 0.01). The temporal generalization maps for visible and invisible cases show that the visible trials display a stronger decoding performance and more generalizable patterns across time, especially across the later time points, suggesting that recurrent processing may aid in the maintenance of representations. We can also see that the difference between visible and invisible trials start to be significant ∼180 ms after target onset which is consistent with the decoding results presented in Figure 3B.

**Figure 4.**
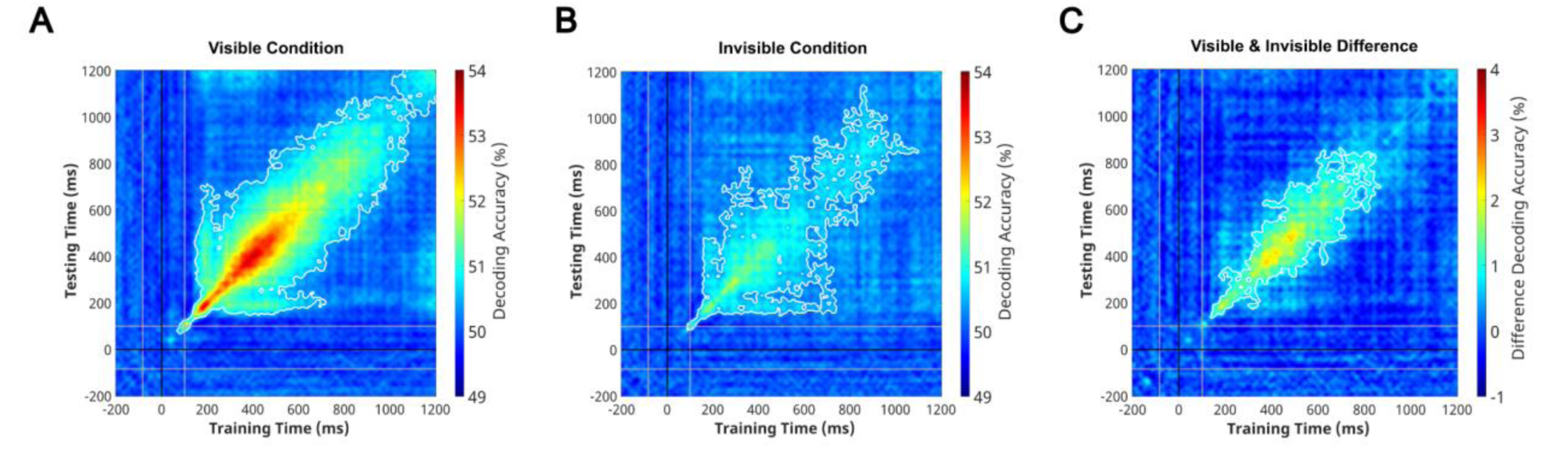
| Temporal Generalization Maps. (A, B, C) Temporal generalization matrices for the visible and invisible conditions and the difference between these two conditions. The white contours show significant decoding values (N=30; one-sided sign permutation tests, cluster defining threshold *p* < 0.01, and corrected significance level *p* < 0.01). The gray lines show the RSVP onset/offset and the black line shows the target onset.

### 4. Gist perception precedes visual identification

In previous sections, we investigated the temporal dynamics of visible and invisible trials separately and compared them to each other. But what if the participant had identified the category correctly, but not the target itself? We separated the trials into the following categories: “both correct” (where both the category and the identity of the target image were reported correctly), “only category correct” (where only the category of the target image – and not the identity – was reported correctly), and “both incorrect” (where both the category and identity of the target image were reported incorrectly). Average behavioral results for each subcategory of images across all participants are shown in Figure 5A. We then calculated the decoding accuracy of the SVM classifier in separating pattern vectors associated with the mentioned cases. Average decoding accuracies across all participants in discriminating “only category correct” trials from the other two types of trials are depicted in Figure 5B. The decoding between “both correct” and “only category correct” isolates the temporal neural trace of image identification (image details perception) which rises significantly above chance ∼190 ms after the target onset. This is later in time compared to only category perception (image gist perception) which is captured by the decoding between “only category correct” and “both incorrect” and rises significantly above chance ∼120 ms post-stimulus.

**Figure 5.**
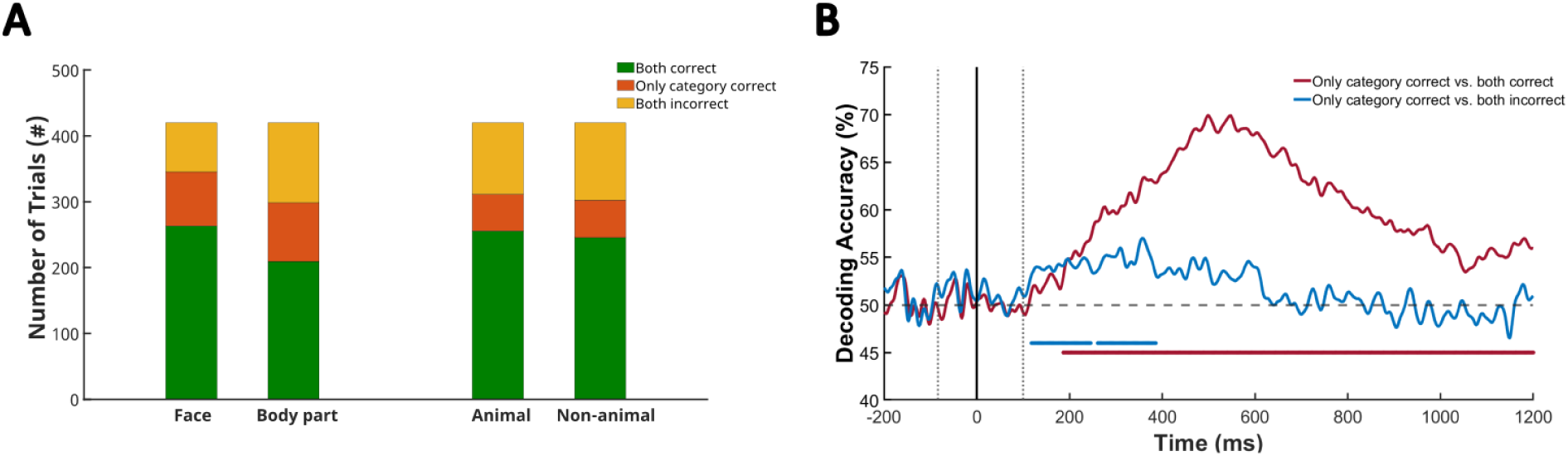
| Categorization vs. Identification. (A) Average behavioral results across all participants. The green bars indicate the condition in which the participant has chosen the both category and identity of the target image correctly, the red bars indicate the condition in which the participant has only chosen the category correctly (and not the identity of the target image), and the yellow bars indicate the condition in which the participant has chosen both the category and the identity of the target image incorrectly. (B) Average decoding accuracies for “only category correct” vs. “both correct” (the red curve) and “only category correct” vs. “both incorrect” (the blue curve). Color-coded bars below the plots show significant time points for each condition. The horizontal dashed line indicates chance level performance. The vertical dotted lines show the RSVP onset/offset and the solid line shows the target onset. (N=30; significant time points were evaluated with one-sided sign permutation tests, cluster defining threshold *p*<0.05, and corrected significance level *p*<0.05)

### 5. The dynamics of recurrent processing are category-dependent

In our experimental design, we incorporated various categories of target images (people category which consisted of images of faces and body parts, and objects category which had images of animals and non-animals). Subsequently, we investigated whether the timing of recurrent processing associated with visibility is dependent on the target category. We probed when and how subcategories are processed differently, reflecting the timing of categorization in the human brain. Figure 6A shows the grand total decoding accuracy separately for people and objects. The grand total curve for the people category has two salient peaks at 105 ms and 183 ms after stimulus onset. These peaks respectively happen 103 ms and 278 ms after the target onset for the objects category. The first peak typically associated with feedforward processing shows a similar timing in both categories, but the discrepancy in the timing of the second peak suggests that recurrent processing may reflect categorical differences. Furthermore, a multidimensional scaling visualization can depict the relations across the neural patterns elicited by the 20 target images for each category (Figure 5B). This visualization shows that for the first peak in both categories, subcategories are not separated from each other; however, for the second peak, images for each subcategory are well separated from the other.

**Figure 6.**
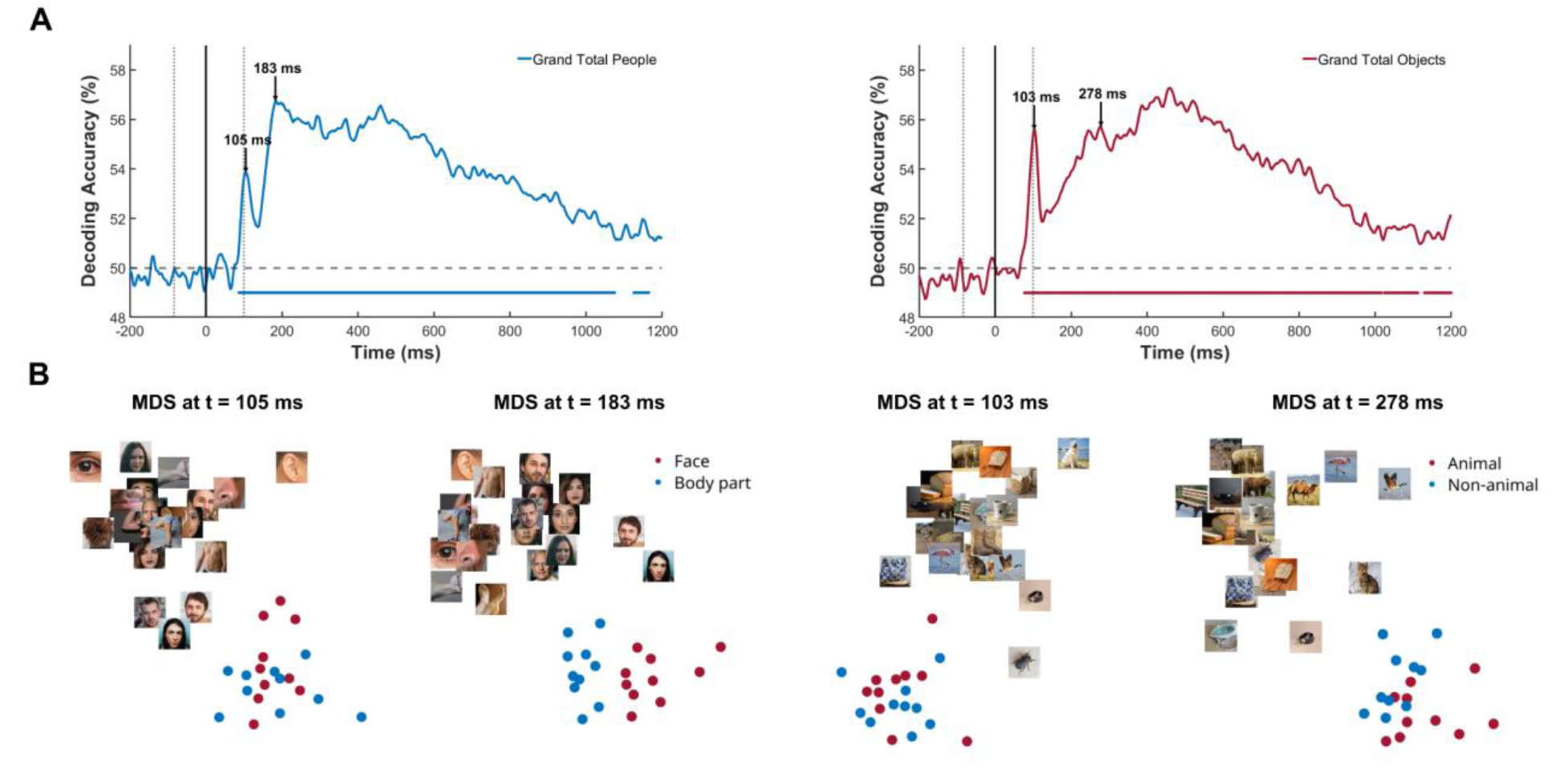
| Categorization in the human brain. (A) Grand total curves for the people and objects categories. The people category consists of images of human faces and human body parts, and objects category includes images of animals and non-animal objects. The dashed lines indicate chance level performance. The dotted lines show the RSVP onset/offset and the solid lines show the target onset. Peak latencies are indicated with arrows. Color-coded bars below the plots show significant time points. (B) MDS plots for the time points of peaks, which are depicted by arrows.

## Discussion

In the present study, we examined the timing and sequence of brain activity involved in recognizing target images correctly (visible) or failing to recognize them (invisible) in the RSVP paradigm. We explored how both initial feedforward and subsequent recurrent neural processes contribute to these visual recognition outcomes. To preserve the attributes of individuals’ daily experiences, we used naturalistic images of people and objects. We separated trials based on participants’ behavioral performance into visible and invisible, and studied the differences between these two behavioral outcomes in EEG data. A multivariate pattern analysis across all conditions showed two salient peaks, at ∼100 ms and ∼200 ms after target onset (Figure 3A).

Evidence from primate and human studies has shown that fast feedforward processing reaches the highest level of the visual hierarchy during the first 100-150 ms post-stimulus onset^14,24,49^. Based on this evidence, the timing of the first peak in the decoding times series reflects fast feedforward processing. However, the second peak happens later in time, indicating more complex recurrent processing. This is also evident in category-specific processing. Within the people and objects categories, the early feedforward decoding peak occurs at a similar latency (∼100 ms). However, the second peak happens at different latencies, which may be an indication of different temporal dynamics and demands for recurrent processing to resolve different categories^23,50^. On the other hand, the multivariate pattern analysis of visible and invisible trials (Figure 3B) shows that these two cases are processed similarly during the early, feedforward stage. This aligns with our ERP findings (see Figure S1) and is in line with previous research, which demonstrated that both visible and invisible trials exhibit comparable amplitudes and topographic distributions in the initial processing stage, represented by the early N/P100 component^34^.

Although subjects were not aware of the target stimuli in the invisible trials, feedforward processing remains undisturbed in line with numerous studies on unconscious processing^24,34,51^. Along with the limitations that feedforward neural network architectures have in object recognition under challenging conditions^33^, this adds to the evidence that the feedforward sweep is not sufficient for the conscious processing of visual objects in challenging conditions. This points to the contribution of other stages of processing in perceiving visual objects in such conditions^5,14,18,52^.

The visible and invisible decoding time series diverge at ∼180 ms after the target onset, with an early peak at ∼200 ms, which is after the time usually associated with the fast initial feedforward processing^14,49,52^. Although the first significant peak in the difference between the visible and invisible time series happens at ∼200 ms, this difference reaches its highest amplitude at ∼400 ms after the stimulus onset.

Fahrenfort et al. (2008) suggested five stages of visual processing in visual tasks. The first stage involves activity in bilateral parietal regions peaking at 121 ms, possibly resulting from feedforward processing. The second stage of visual processing involves a more posterior occipital activity and peaks at 160 ms. This stage may show a combination of feedforward and feedback activity within the early visual cortex. The third stage of processing peaks at ∼200 ms in the occipitoparietal and centrofrontal regions and may reflect perceptual-level attentional selection. The fourth stage involves activity in more posterior occipital regions and peaks at 246 ms, which is possibly related to recurrent activity enabling conscious access. Lastly, the final stage of processing peaks at ∼350-400 ms in occipitoparietal brain regions, reflecting recurrent processing within and/or between these areas. Within this stage, they observed a posterior-parietal component, commonly referred to as the P3 or P300, in the ERP waveform. This P3 component has been linked to psychological factors such as working memory and attention^52^.

Thus, while visible and invisible cases reflect similar feedforward processing, there is an important difference in whether they reach visual awareness and trigger attentional processes (see Figure 3B). The temporal dynamics observed in our data suggest it is likely that in the invisible cases, recurrent connections are not triggered and this inhibits their further processing into visual awareness. This is supported by the observation of the stronger performance of EEG decoding in the visible trials compared to the invisible trials, as well as the maintenance of later-stage activity in the visible cases, potentially reflecting the presence of attention.

On the other hand, the temporal generalization results qualitatively show different maps for the visible and invisible trials. We can see that the visible trials depict a broader map with higher decoding accuracy in comparison to the invisible trials. This is true, especially in the later time points which can be associated with recurrent processing or consciousness. In the visible trials, neuronal activity seems to be more persistent during these later time points (especially between 200 and 500 ms after the target onset) and this persistent activity could maintain the results of the recurrent processing for later use –for instance reaching consciousness or making decisions^45^.

We also disentangled target categorization from target identification. Previous psychophysical studies in humans have suggested that one can get the gist of complex visual images within about 150 ms, even when the stimulus itself is presented as briefly as 10 ms, but it generally takes longer to identify individual objects^37,39^. Moreover, it may take even longer for a fuller semantic understanding of the image to emerge and be encoded in short-term memory^53^. We assumed that when participants get the gist of the image, they can recognize the category of the image; however, correct identification only happens when they report the target accurately. The decoding time series of “only category correct” vs. “both correct” (the red curve in Figure 5B) shows when these two conditions start to diverge from each other at ∼190 ms. We believe that categorization occurs in both conditions (”both correct” and “only category correct”), but the correct identification of the image only occurs in the “both correct” condition, thus causing the different decoding time courses. Similarly, the difference between “only category correct” and “both incorrect” conditions (the blue curve in Figure 5B) may be associated with categorization in the brain. The decoding accuracy of “only category correct” vs. “both incorrect”, which shows the presence of categorization, rises significantly above chance about 120 ms post-stimulus. This timing suggests that the gist of a complex image can be perceived solely through feedforward processing of the visual signal, as shown by previous studies^2,38,54^. However, when it comes to identification, the decoding accuracy of “both correct” vs. “only category correct” doesn’t rise significantly above chance until ∼190 ms after the target onset. This timing is likely to reflect the recurrent processing involved in the identification of the target image.

The present study highlights the use of EEG for shedding light on challenging questions about perceptual and cognitive processing in the human brain. Here we see a clear benefit to the high temporal resolution of EEG in differentiating processing mechanisms, such as those underlying visible versus invisible images, in identification versus discrimination, and in gist versus object recognition. Future studies could employ complementary neuroimaging techniques like functional Magnetic Resonance Imaging (fMRI) to leverage higher spatial resolution in localizing the sources of feedforward and recurrent processes involved in visual processing. Furthermore, a deeper investigation of different stimulus categories and the sources of category-specific feedback processing using fMRI may elucidate categorical processing in the brain.

## Conclusion

This study was conducted with the aim of explaining the difference in the processing of visible and invisible stimuli when they are under challenging visibility conditions. The results indicate that visible and invisible cases share a similar temporal dynamic during the first stages of visual processing, which is reflected by the initial fast feedforward processing; however, during the later stages of processing, there is a significant difference between these two cases. This difference might be related to the recurrent processing in the human brain, which is not completely triggered in the invisible cases and as a result, the human brain will not perceive target stimuli consciously.

## Methods

### 1. Participants

32 neurologically healthy adult participants participated in this experiment. However, only the data from 30 participants (16 female, mean age: 24.13, sd 4.78 years) were used in the study as two of the participants were excluded from the experiment, one due to their low performance which was around the chance level, and the other one because of their very high performance in comparison to other participants which made it an outlier. Participants were able to read and write in English, were right-handed, had normal/corrected-to-normal vision, and had no history of neurological diseases. Written consent was obtained from all participants. The study was approved by The Health Sciences Research Ethics Board at Western University, and all participants were compensated for their time.

### 2. Stimulus set and experimental design

The stimulus set consisted of 20 color images of people (with 10 pictures of human faces and 10 pictures of human body parts) and 20 color images of objects (with 10 pictures of animal objects and 10 pictures of non-animal objects). Images were chosen from the THINGS and the Real and Fake Face Detection databases^55,56^. We used the Natural Image Statistical Toolbox^57^ to control for potential confounds in low-level visual features such as color and spatial frequency (Tables S1, S2, S3, S4).

Participants viewed a rapid series of 11 images, each presented for 17 ms at the center of the screen (7.7° visual angle) against a gray background using Psychtoolbox in MATLAB^58,59^. The middle (sixth) image was the target, and the remaining ten images were scrambled images generated using our target images (by dividing each image in the stimulus set into 900 patches and shuffling these patches randomly; see Figure 1). Each participant completed 6 runs for each category (people and objects), and each run lasted ∼6 minutes. There were 140 trials in each run. Each trial started with a 500 ms fixation period, followed by the image series. At the end of each trial, after a delay of 800–1000 ms (uniformly distributed), participants were instructed to choose the target image among four options. For the people category, the options consisted of two face and two body part options, including the target image and three randomly chosen lure options; similarly, for the objects category, the options consisted of two animal and two non-animal object options, including the target image and three randomly chosen lure options. Participants had 1.2 seconds to respond via a button press. Every target image was presented 7 times in random order in each run and 42 times over the duration of the experiment.

Prior to the start of the experiment, participants completed a ∼5-minute practice block to familiarize them with the task. Next, they were cued to the stimulus set by presenting each target image for 2 seconds.

### 3. EEG data recording and preprocessing

EEG data were recorded using a 64-channel Biosemi system at a sampling rate of 2048 Hz. All channels were referenced to the CMS channel (Common Mode Sensor). The data were preprocessed using the Brainstorm Toolbox in MATLAB^60^. A bandpass filter with cut-off frequencies of 0.5 and 30 Hz was applied to the data to remove slow drifts and high-frequency artifacts including powerline noise. The blink artifacts were removed using the ICA method. To speed up the computational process, EEG signals were down-sampled at a rate of 512 Hz.

### 4. Multivariate Pattern Analysis of EEG data

Multivariate pattern analysis of EEG data followed methods set out in^14,61^. EEG data was first segmented by extracting EEG signals from –200 to 1200 ms with respect to the target onset. To investigate how target images diverge from each other in the human brain over time, we formed representational dissimilarity matrices (RDMs) for each participant. Each column and row of these matrices are indexed with our target images, and the cells contain the dissimilarities between all pairs of images as assessed using the accuracy of a linear SVM that discriminates the neural responses of each pair of target images. To reduce computational complexity, we divided our trials into five folds and averaged the trials in each fold to create five pseudotrials. The performance of the classifier was then recorded with the leave-one-out cross-validation method. At each iteration, one of the pseudotrials was used for testing and the rest was fed to the classifier as the training set. We averaged all elements of the RDM to form the grand total decoding series for each participant and then averaged these over all participants (Figure S2).

Visible and invisible cases were identified as trials in which the participant identified the target image correctly (visible trial), or incorrectly/ did not report the target at all (invisible trial). For each case, we used a linear SVM to discriminate between EEG pattern vectors for each pair of images.

To overcome the imbalance in the number of trials for visible and invisible image conditions, we used a bootstrapping technique which subsampled EEG trials for each label with the number of trials in the minority class, repeated this 100 times, and averaged the decoding accuracies of these 100 repetitions. Conditions with fewer than 5 trials were excluded from the analysis. For each participant, we averaged all the elements in visible and invisible RDMs to generate the visible and invisible decoding time series, and at the last step, we averaged these decoding time series over all participants.

### 5. Temporal generalization analysis

Neuronal activity, reflecting different neural processes, can either remain constant over time or change rapidly. Undoubtedly, if the neuronal activity is persistent over some time points, EEG signals should be similar across that time interval^45^. To examine whether the neural representations discriminating between visible and invisible cases are persistent for extended times along the ventral visual pathway, we used the temporal generalization approach^46^. To perform this analysis, EEG pattern vectors were extracted for all time points and all conditions similar to the previous section (see Multivariate Pattern Analysis for EEG data). We trained a linear SVM classifier to pairwise decode EEG data for different target images on time point (t) and then tested our classifier using EEG data at all other time points (t’). Performance of the classifier for all image pairs was averaged and formed the element (t,t’) in the temporal generalization matrix. This was repeated for all pairs of time points to form temporal generalization matrices for the visible and invisible cases. As this process was time-consuming and computationally complex, the EEG signals were down-sampled at 256 Hz exclusively for this analysis (Figure S3). Temporal generalization maps with stronger activities and broader off-diagonal maps across future time points indicate more persistent neuronal activities.

### 6. Categorization vs. target identification in the human brain

To discriminate the temporal dynamics of object identification and object categorization, we separated all the trials in which participants reported the target correctly (which means they have identified the target image) and we categorized them as “both correct”. The trials in which participants only chose the category of the target correctly (which means they categorized the object, but did not correctly identify the image) were categorized as “only category correct”. Lastly, the trials in which both category and identity were reported incorrectly, were categorized as “both incorrect”. This resulted in three conditions. Similar to the analysis in section 4, we used a linear SVM to discriminate between EEG pattern vectors for each pair of the three cases. As the number of trials for the “both correct” case was higher in comparison to the other two conditions, we performed bootstrapping to subsample the trials in this case (see section 4 for detailed information about bootstrapping).

### 7. Multidimensional scaling (MDS)

The EEG RDMs capture the relationships between neural patterns related to the 40 target images. In order to visualize the complex patterns of these RDMs we used multidimensional scaling (MDS), an unsupervised approach to visualizing similarity among diverse conditions within a distance matrix. MDS maps the data to a two-dimensional space where similar images are clustered in close proximity, while dissimilar images are positioned farther apart^62,63^. The MDS plots were generated using the *mdscale* built-in function in MATLAB. This function performs nonmetric multidimensional scaling on the n-by-n dissimilarity matrix.

### 8. Statistical Testing

To correct for multiple comparisons across time points, we performed permutation tests for cluster-size inference. We used cluster-size inference with 10,000 permutations, and a cluster-defining threshold of *p* < 0.01 for all grand total decoding time series, visible and invisible decoding time series, the difference time series, and all temporal generalization maps, and a cluster-defining threshold of *p* < 0.05 for “only category correct” vs. “both correct” and “only category correct” vs. “both incorrect” time series. The null hypothesis was 50% chance level for the classification time series and 0 for the difference decoding time series.

## Acknowledgement

The authors would like to thank Diana Dima for helpful comments on the manuscript and Mansoure Jahani and Ali Tafakkor for helpful discussions. This study was supported by the Canada First Research Excellence Fund (CFREF) through a BrainsCAN grant to Y.M., and a Vector Institute Research Grant to Y.M., and S.C.M. Also, S.C.M. received a graduate student scholarship from CFREF BrainsCAN.

