## Supplementary Information for "Unveiling the neural dynamics of conscious perception in rapid object recognition"

### ERP comparison for visible and invisible cases

A large body of studies has studied the different stages of processing in visible and invisible cases through ERPs[1,2]. To add more reliability to our data, we completed an ERP analysis comparing the ERP results for visible and invisible cases to those in other studies. To do so, we constructed three electrode clusters: Frontal (FP1, FP2, F3, Fz, and F4), Central (FC1, FC2, C3, Cz, and C4), and Parietal (CP3, CP4, P3, Pz, and P4). We averaged the ERPs for all the electrodes in each cluster and compared visible and invisible trials to each other (Figure S1A, S1B, and S1C). Results show that these cases share similar amplitudes at early time points (N/P100), consistent with a similar initial feedforward sweep in processing for both trial types. The two cases begin to diverge ~330 ms after the target onset in all three clusters, suggesting different later processing in the two trial types. Note, however, that the multivariate pattern analysis of visible and invisible cases showed a significant divergence much earlier, beginning ~180 ms after the target onset. This discrepancy in timing suggests the multivariate pattern analysis is more sensitive to changes between conditions that are not apparent in averaged ERPs.

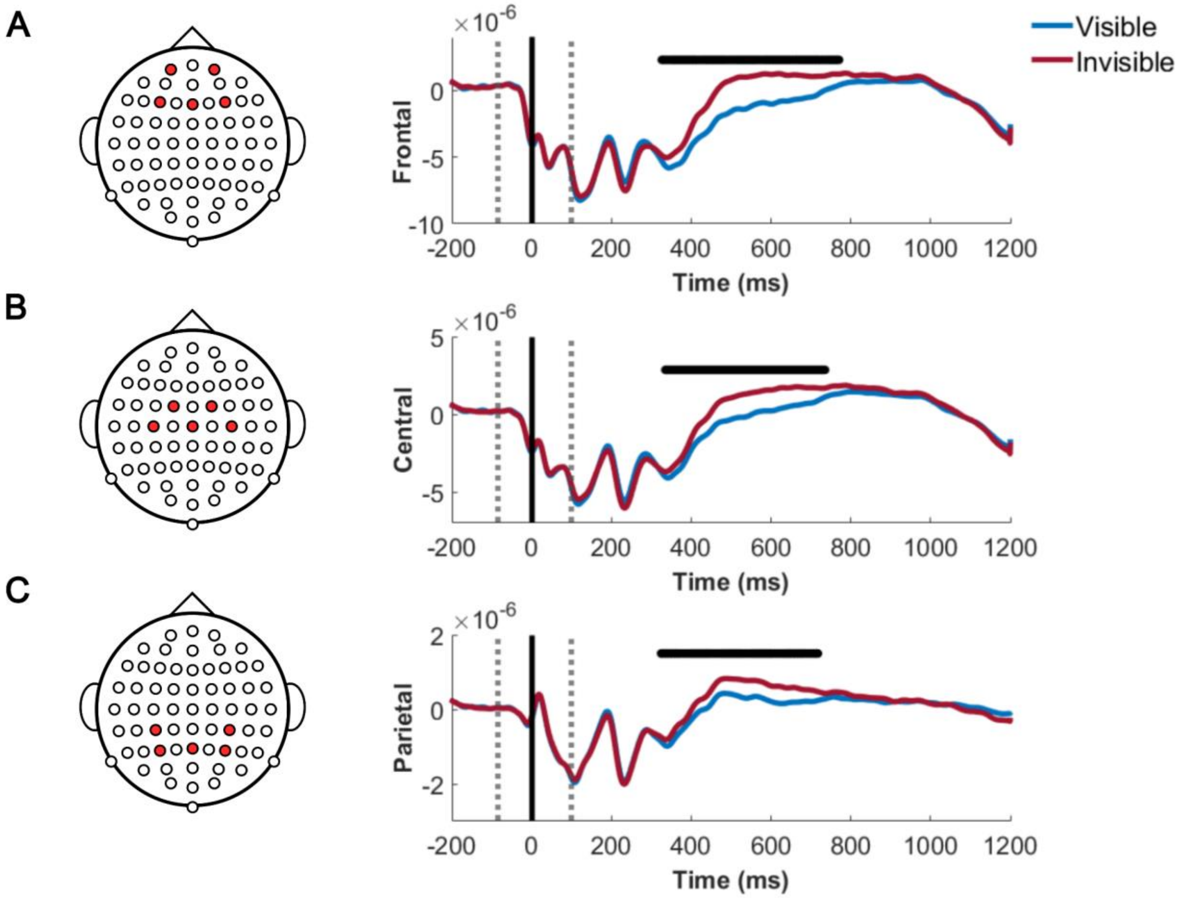


**Figure S1 | ERPs of Frontal, Central, and Parietal clusters in visible and invisible conditions.**

(A) ERPs for visible and invisible conditions in the Frontal cluster. FP1, FP2, F3, Fz, and F4 electrodes are averaged for each condition. (B) ERPs for visible and invisible conditions in the Central cluster. FC1, FC2, C3, Cz, and C4 electrodes are averaged for each condition. (C) ERPs for visible and invisible conditions in the Parietal cluster. CP3, CP4, P3, Pz, and P4 electrodes are averaged for each condition.

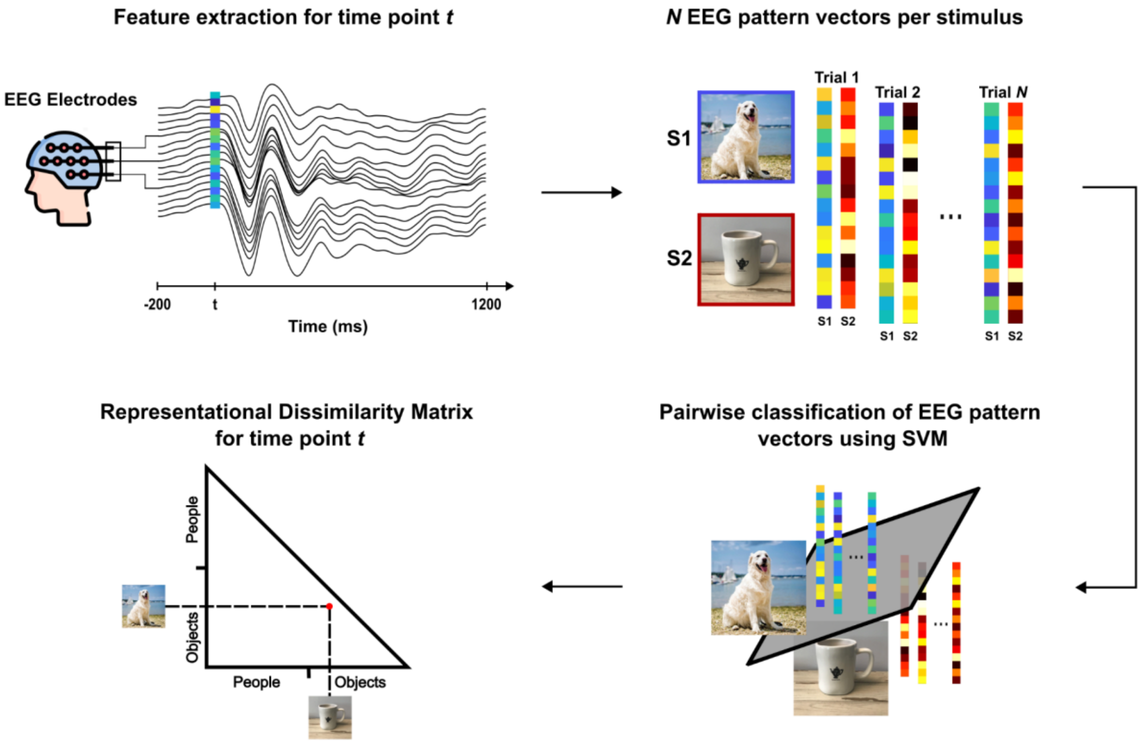


**Figure S2 | Multivariate Pattern Analysis.**

EEG pattern vectors were extracted from trials at time point t. A support vector machine classifier was used to pairwise classify EEG pattern vectors for target images. Representational Dissimilarity Matrices (RDMs) were formed for each time point using classifier performance.

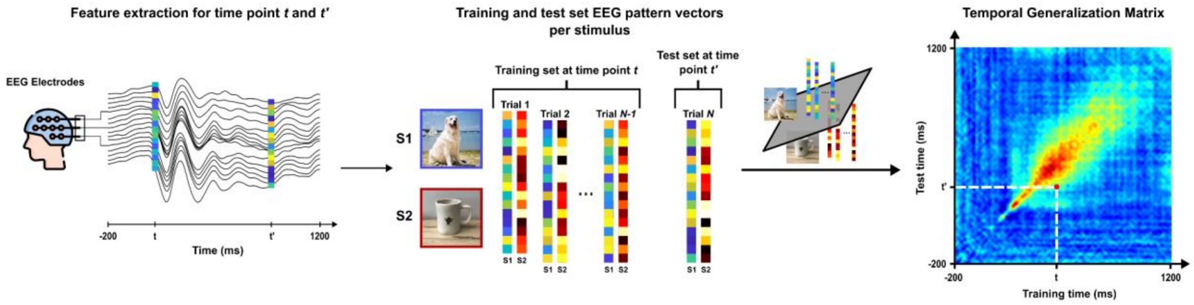


**Figure S3 | Temporal Generalization approach.**

EEG pattern vectors were extracted from trials at time points t and t'. A support vector machine classifier was trained on data at time point t and then tested on data at time point t'. Temporal generalization decoding matrices were formed for different conditions using the classifier performance.

**Tabel S1 | Comparison of the spatial frequency information between faces and body parts**

| | Spectral Energy Level (Spatial Frequency) | | | | |
|---|---|---|---|---|---|
| | **10%** | **30%** | **50%** | **70%** | **90%** |
| $t - value$ | -0.4932 | 0.4932 | 0.6061 | 0.4845 | -0.4845 |
| $p - value$ | 0.6278 | 0.6278 | 0.5520 | 0.6338 | 0.6333 |

**Tabel S2 | Comparison of color distribution (RGB and Lab) between faces and body parts**

| | RGB Space | | | Lab Space (Brightness & Contrast) | | |
|---|---|---|---|---|---|---|
| | **R** | **G** | **B** | **L** | **a** | **b** |
| $t - value$ | -1.7719 | -1.3319 | -0.9577 | -2.0997 | -1.3910 | -1.5204 |
| $p - value$ | 0.0933 | 0.1995 | 0.3509 | 0.0501 | 0.1812 | 0.1458 |

**Tabel S3 | Comparison of the spatial frequency information between Animals and Non-animals**

| | Spectral Energy Level (Spatial Frequency) | | | | |
|---|---|---|---|---|---|
| | **10%** | **30%** | **50%** | **70%** | **90%** |
| $t - value$ | -1.5000 | -0.7767 | -0.3416 | -0.5829 | -0.8938 |
| $p - value$ | 0.1510 | 0.4474 | 0.7366 | 0.5672 | 0.3832 |

**Tabel S4 | Comparison of color distribution (RGB and Lab) between Animals and Non-animals**

| | RGB Space | | | Lab Space (Brightness & Contrast) | | |
|---|---|---|---|---|---|---|
| | **R** | **G** | **B** | **L** | **a** | **b** |
| $t - value$ | -0.3489 | 0.5244 | 0.8129 | 0.5145 | -1.2677 | -0.7683 |
| $p - value$ | 0.7312 | 0.6064 | 0.4269 | 0.6131 | 0.2211 | 0.4523 |
